# The effect of morphine on rat microglial phagocytic activity: an in vitro study of brain region-, plating density-, sex-, morphine concentration-, and receptor-dependency

**DOI:** 10.1101/2022.10.03.510683

**Authors:** David N. King’uyu, Lily Nti-Kyemereh, Jesse L. Bonin, Paul J. Feustel, Michelle Tram, Katherine C. MacNamara, Ashley M. Kopec

## Abstract

Opioids have long been used for clinical pain management, but also have addictive properties that have contributed to the ongoing opioid epidemic. While opioid activation of opioid receptors is well known to contribute to reward and reinforcement, data now also suggest that opioid activation of immune signaling via toll-like receptor 4 (TLR4) may also play a role in addiction-like processes. TLR4 expression is enriched in immune cells, and in the nervous system is primarily expressed in microglia. Microglial phagocytosis is important for developmental, homeostatic, and pathological processes. To examine how morphine impacts microglial phagocytosis, we isolated microglia from adult male and female rat cortex and striatum and plated them *in vitro* at 10,000 (10K) or 50,000 cells/well densities. Microglia were incubated with neutral fluorescent microbeads to stimulate phagocytosis in the presence of one of four morphine concentrations. We found that the brain region from which microglia are isolated and plating density, but not morphine concentration, impact cell survival *in vitro*. We found that 10^-12^M morphine, but not higher concentrations, increases phagocytosis in striatal microglia *in vitro* independent of sex and plating density, while 10^-12^M morphine increased phagocytosis in cortical microglia *in vitro* independent of sex, but contingent on plating density. Finally, we demonstrate that the effect of 10^-12^M morphine in striatal microglia plated at 10K density is mediated via TLR4, and not µORs. Overall, our data suggest that in rats, a morphine-TLR4 signaling pathway increases phagocytic activity in microglia independent of sex. This may be useful information for better understanding the possible neural outcomes associated with morphine exposures.

## INTRODUCTION

Opioids have long been used for clinical pain management, but also have addictive properties (Strang et al., 2020; Volkow et al., 2019; Yoo et al., 2022) that have contributed to the ongoing opioid epidemic (Stoicea et al., 2019; Terplan, 2017). The opioid morphine binds to both opioid receptors (such as mu-opioid receptor, µOR) which are present on several different cell types, and toll-like receptor 4 (TLR4), which is enriched on immune cells (X. Wang et al., 2012; Xie et al., 2017). Opioid receptors are coupled to Gi/o proteins that inhibit adenylate cyclase, decrease cyclic adenosine monophosphate levels, and recruit β-arrestin which activates p38, AKT and ERK pathways (Al-Hasani & Bruchas, 2011; Fu et al., 2014; Valentino & Volkow, 2018). TLR4 is coupled to MyD88 and TRIF which stimulate the translocation of NF-κB to the nucleus leading to production of inflammatory cytokines and type I interferons (Lu et al., 2008; Rahimifard et al., 2017). Functionally, in peripheral immune cells opioid receptor activation induces synthesis and release of opioid peptides (Boué et al., 2011; Schäfer et al., 1994), while TLR4 activation generally stimulates pro-inflammatory and phagocytic clearance activity (Ninkovic et al., 2016; Schaub et al., 2004). Phagocytic activity is a hallmark of immune macrophages that serves to clear damage, debris, and infection (Andoh & Koyama, 2021). While opioid activation of opioid receptors is well known to contribute to reward and reinforcement, recent data suggest that opioid activation of TLR4 also plays a role in addiction-like processes (Hutchinson et al., 2012; Wu & Li, 2020; Zare et al., 2023) via expression of pro-inflammatory cytokines TNFα and IL-1β (Pascual et al., 2012; Rizzo et al., 2018; Xiaohui Wang et al., 2012). In the nervous system, TLR4 expression is primarily expressed in microglia, the resident immune cells of the central nervous system (Zhang et al., 2014; Zhang et al., 2016). Microglial phagocytosis is important for developmental, homeostatic, and pathological processes (Fu et al., 2014; Paolicelli et al., 2022). Thus, understanding the diverse actions of opioids via these different receptors, is important to develop strategies to mitigate their negative consequences.

In peripheral immune cells, morphine reproducibly decreases immune activation and phagocytosis *in vivo* and *in vitro* (Eisenstein, 2019; Ninkovic et al., 2016; Ninković & Roy, 2012; Rojavin et al., 1993) dependent on concentration (Borman et al., 2009) and duration of exposure (Pagán et al., 2001). Unlike peripheral immune cells, the effect of morphine on microglia is less well studied. Morphine has been reported to both increase and decrease microglial phagocytosis (Lipovsky et al., 1998; Peterson et al., 1995; Ryu et al., 2018; G. Sowa et al., 1997). There is no consensus on the modulatory effect of morphine on microglia phagocytosis currently. Several different factors contribute to microglial phagocytic activity, including sex, brain region, and regional microglial density. For example, during neonatal development microglia from female mice are more phagocytically active in the hippocampus (Nelson et al., 2017) and mature faster (Hanamsagar et al., 2017) than microglia from male mice. Morphine increases the number of activated microglia in the female rat periaqueductal gray compared to male rats which correlates to morphine analgesia (Doyle et al., 2017). Morphine also leads to sex specific microglia polarization in the periaqueductal gray (Doyle et al., 2017).

There also appears to be heterogeneity in the transcriptional profile of cortical and striatal microglia (De Biase et al., 2017; Grabert et al., 2016). Deep single cell RNA sequencing reveals that microglia transcriptomes are heterogenous (Li et al., 2019) indicating features that may underlie microglia variability across brain regions (H. S. Yang et al., 2021). This heterogeneity suggests that morphine results in different responses depending on the transcriptional identity of the microglia which are spatially regulated (Coffey et al., 2022) potentially modulating the normal tendency to phagocytose. Finally, the *in vivo* density of microglia varies depending on brain region with the highest density in the substantia nigra (Lawson et al., 1990; Yang et al., 2013). A higher number of microglia in a region correlates with increased maladaptive neuroinflammation and/or microglial ‘states’ (Bollinger et al., 2017; Furube et al., 2018; Tynan et al., 2010). Morphine decreases the number of microglia in the substantia nigra (Jokinen et al., 2018).

We hypothesized that morphine would modulate phagocytic activity in microglia *in vitro*, but in a culture density-, concentration-, region-, and sex-dependent manner. Specifically, we predicted microglia would be more phagocytically active, and thus less responsive to the suppressive effects of morphine if they are (1) plated at higher densities relative to lower densities, (2) from striatum relative to cortex, and (3) from females relative to males. To test this, we isolated microglia from adult male and female rat cortex and striatum and plated them *in vitro* at a density of 10,000 or 50,000 cells (10K or 50K hereafter). Microglia were incubated with neutral fluorescent microbeads to stimulate phagocytosis in the presence of one of four morphine concentrations (**Fig. 1**). Unexpectedly, we found that we could not directly compare cortical and striatal microglia, nor the two culture densities, as region and density, but not morphine concentration, impacted cell survival over the course of the experiment. When we separated region and density conditions, we observed that 10^-12^M morphine, our lowest tested concentration, increased phagocytic activity in 10K striatal, 50K striatal, and 50K cortical cells, but not 10K cortical cells. The activating effect of 10^-12^M morphine on 10K striatal, but not 10K cortical cells, was replicated in separate studies, and we further provide evidence that TLR4s, and not µORs is the primary receptor mediating this effect. Overall, our data suggest that a morphine-TLR4 signaling pathway increases phagocytic activity in microglia independent of sex, which is useful information for better understanding the possible neural outcomes associated with morphine exposures.

**Figure 1:**
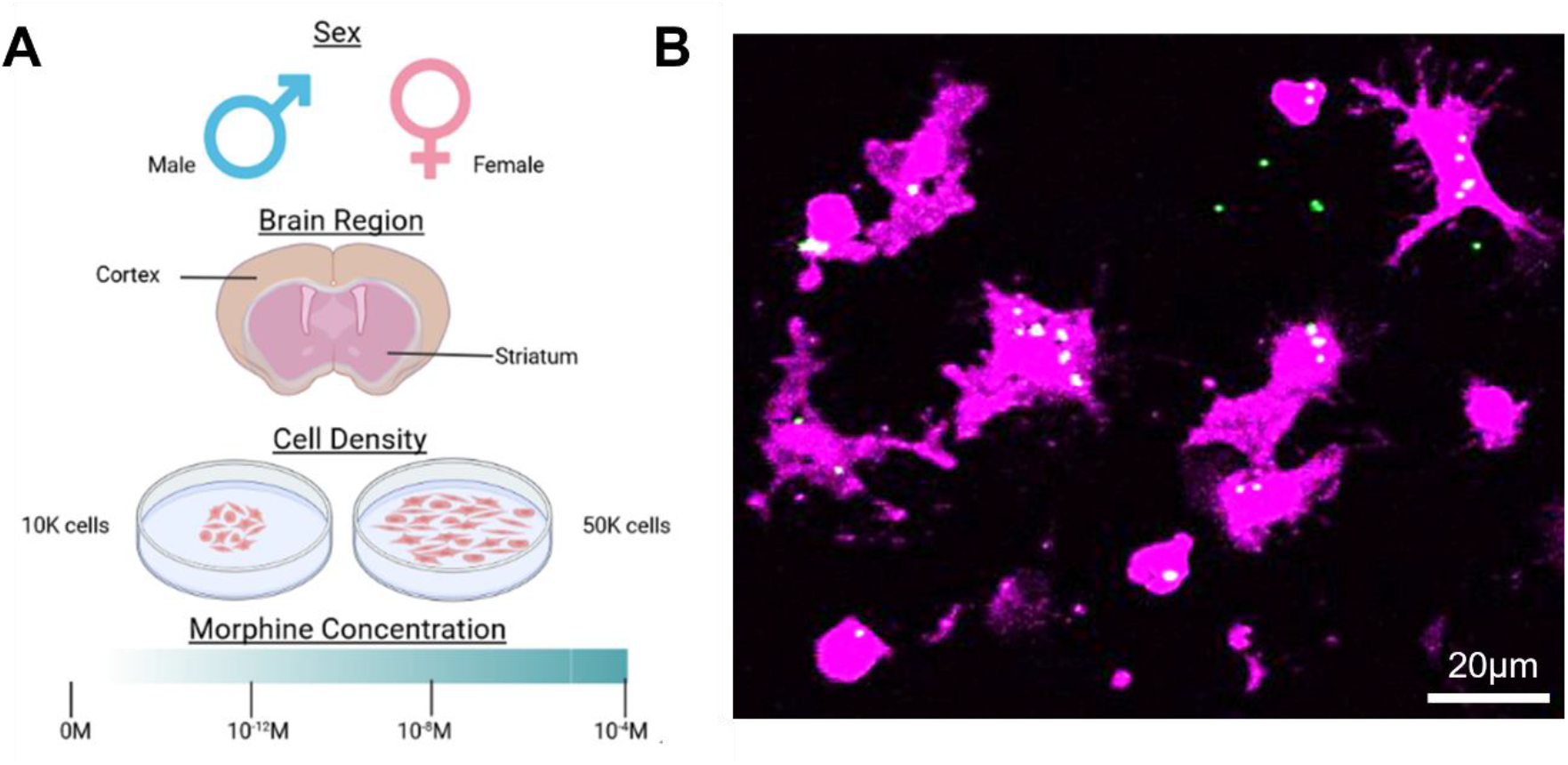
Experimental Design. **(A)** The experimental design included sex (male vs. female), brain region (cortex vs. striatum), density (10,000 microglia/well (10K microglia) vs. 50,000 microglia/well (50K microglia), and morphine concentration (0, 10^-12^, 10^-8^, and 10^-4^ M) as independent variables. Created with Biorender.com. **(B)** Representative image of Iba1+ microglia (magenta), fluorescent microbeads (green), and colocalization of microglia and fluorescent microbeads (white) which we refer to as phagocytosis. Image is representative of all conditions.

## MATERIALS AND METHODS

### Animals

Young adult (postnatal day (P)70-P90) male and female Sprague Dawley rats were purchased (Envigo) and group-housed with ad libitum access to food and water for at least 2 weeks. Conditions were kept at 23°C on a 12:12 light:dark cycle (lights on at 07:00) and cages were changed twice a week. Experiments and animal care were approved by the Institutional Animal Care and Use Committee at Albany Medical College.

### Tissue collection and microglial isolations

Animals were euthanized by CO_2_ anesthesia and exsanguination. Saline perfusion was performed before animals were decapitated and brains extracted. Each brain had cortex and striatum regions grossly dissected and separated for mechanical and enzymatic breakdown. Brain regions were minced using a razor into a gelatinous consistency while on a petri dish resting on wet ice. Minced brain tissue underwent enzymatic dissociation using the Neural Tissue Dissociation Kit (P) protocol (Miltenyi Biotec, 130-092-628) to isolate neural cells. Neural cells were incubated with CD11b+ magnetic microbeads (Miltenyi Biotec, CD11b/c Microbeads, rat, 130-105-634) and CD11b+ cells positively selected through a magnetic column to enrich for microglial cells. We will refer to CD11b+ cells hereafter as microglia. Microglia comprised on average 28.5% of live cells in pre-isolation samples, and 72.3% of live cells after CD11b+ isolation **(Supplemental Fig. 1).**

### Culture plating

Microglia were incubated in media (Dubelco’s Modified Eagle’s Medium with 1µM forskolin and 1% N2 media, 1% L-glutamine, 1% sodium pyruvate, 1% Pen-Strep by volume (v/v)) at 37 °C/5% CO_2_ in chambered slides (Thermo Fisher Scientific Nunc Lab-Tek II CC^2^ Chamber Slide, Z734853) at either 10K microglia (10,000 microglia/well or 50,000 microglia/mL) and 50K microglia (50,000 microglia/well or 250,000 microglia/mL) density. Microglia were counted using trypan blue (live-dead assay) on an automated cell counter (TC20 Automated Cell counter, Biorad, 508BR08836).

### Cell survival analysis

Isolated microglia were counted for pre-incubation survival. 10K and 50K microglia were plated in a 96 well plate and incubated at 37⁰C for 2hrs in 200µL media. Morphine sulfate reconstituted in 100µL was added at 10^-12^ M, 10^-8^ M and 10^-4^ M to samples and incubated for 90 mins (final volume 300µL). Microglia were counted for post-incubation survival.

### Flow cytometry analysis of enrichment

Neural cells were divided into two fractions which we refer to as either pre-enriched or enriched. Enriched cells underwent magnetic microbead separation to isolate CD11b+ cells. CD11b-enriched and pre-enriched cells were stained with Zombie Red viability dye (BioLegend Cat. No. 423110) and the surface markers CD11b/c (Thermofisher Scientific Cat. No. 12-011082) and CD45 (Thermofisher Scientific Cat. No. 17-0461-82). Samples were acquired on a Northern Lights Spectral Flow Cytometer (Cytek Biosciences, California, US). Data were analyzed using FlowJo (BD, NJ, US) and statistical analysis was performed using Graphpad Prism 9.5.0 (paired *t*-test. p<0.05 was defined as statistically significant.

### Morphine-fluorescent microbead addition

Microglia were permitted to adhere to the chambered slides for 2hrs in 200µL media/well. Fluorescent microbeads (Polysciences Inc., Fluoresbrite Carboxylate YG 1.0-micron microspheres, 15702-10) and morphine sulfate (Mallinckrodt, H11662) at 0 Molar (M), 10^-12^ M, 10^-8^ M and 10^-4^ M were reconstituted in 100µL fresh media/well and was added to seeded microglia (final volume 300µL) simultaneously. This range of morphine concentration is similar the range of concentrations previously used *in vitro* to study microglia phagocytosis ∼10^-6^ – 10^-20^ M (Lipovsky et al., 1998; Peterson et al., 1995; Ryu et al., 2018; G. Sowa et al., 1997). The total amount of microbeads per well is ∼1.0 x 10^8^ microbeads/well. Microglia were incubated at 37⁰C for an additional 90 mins, which we have published is sufficient for phagocytic activity to occur (Kopec et al., 2018).

### Agonist-antagonist pilot assays

Isolated microglia were permitted to adhere to the chambered slides for 1.5hrs in 100 µL media/well. Naltrexone hydrochloride (Sigma-Aldrich, Cat. No. N3136) at 0, 50, 100 and 200 µM or *Rhodobacter sphaeroides* lipopolysaccharide (RS-LPS, InvivoGen, Cat. No. tlrl-prslps) at 0, 1, 5, 10 µg/mL were reconstituted in 100 µL fresh media/well and were added to seeded microglia (volume 200 µL) for 30 mins. (D-Ala(2)-mephe(4)-gly-ol(5))enkephalin (DAMGO, Sigma-Aldrich, Cat. No. E7384) at 50 µM or lipopolysaccharide (LPS, Sigma-Aldrich, Cat. No. L4391) at 1 µg/mL was reconstituted with fluorescent microbeads in 100 µL fresh media/well and were added to naltrexone and RS-LPS pre-incubated microglia respectively (final volume 300 µL) for 1.5hrs.

### Inhibitor-morphine-bead addition

Isolated microglia were permitted to adhere to the chambered slides for 1.5hrs in 100µL media/well. Naltrexone at 100µM or *Rhodobacter sphaeroides* lipopolysaccharide (RS-LPS) at 1µg/mL were reconstituted in 100µL fresh media/well and were added to seeded microglia (volume 200µL) for 30mins. Fluorescent microbeads and morphine sulfate at 10^-12^ M were reconstituted in 100µL fresh media/well and were added to naltrexone and RS-LPS pre-incubated microglia (final volume 300µL) for 1.5hrs.

### Immunocytochemistry

Microglia were rinsed with PBS (2x washes, 300µL/well), then fixed with paraformaldehyde (300 µL/well PFA; 4%) for 20mins. Microglia were permeabilized and blocked using 0.3% Triton x-100 and 10% donkey serum in PBS (v/v), and incubated with primary antibody anti-rabbit Iba1 (1:500; Wako; 019-19741) overnight at 4⁰C. The following day, microglia were rinsed with PBS (3x washes, 300µL/well) and incubated with secondary antibody 1:500 Alexa Fluor donkey anti-rabbit 568 (ThermoFisher, A21202) at room temperature for 2hrs in the dark. Microglia were rinsed with PBS (3x washes, 300µL/well) and chambered slide well walls were removed. Glass slides were dried overnight in the dark. Slides were coverslipped using mounting agent (Invitrogen, Prolong Glass Antifade Mountant, P36984) the following day.

### Imaging and Data Acquisition

2-6 images per condition were captured on a ZEISS LSM 880 confocal microscope at 20x magnification to capture at least 20 Iba1+ structures per sample. The imager was blinded to the animal, region, and sex of each sample. Morphine concentration and density could not be blinded due to uniform plating maps to reduce experimental error. 20µm slice z-stacks, 0.5µm step size, were taken of Texas Red (Iba1+ microglia) and GFP (microbeads) channels. Z-stacks were maximum projected into 2D images with ImageJ (National Institutes of Health; Bethesda, MD). Channels were separated and the microglia channel was manually outlined in the absence of microbead signal. Outlines were drawn by experimenters blinded to the experimental conditions. After all microglia in the image were outlined, cell outlines were overlayed onto the microbead channel and mean gray value per Iba1+ area (µm) was measured. Some Iba1+ structures contained many microbeads, some contained none. A single average value of all GFP+ pixels within the bounds of the Iba1+ areas was used for each condition. A representative image of the phagocytic diversity can be found in **Fig. 1B**.

### Statistical analysis

The endpoint that was statistically analyzed was the amount of phagocytosed beads measured as bead fluorescent intensity within microglia, which represents microbead internalization. Outliers were identified with log transformed data due to heteroskedasticity in Graphpad 9.4.1 using the ROUT method (Q=1%); a single outlier was removed from the data presented in Fig. 4, which is indicated in the figure legend. Statistics were analyzed with SPSS 28.010 or GraphPad 9.4.1 with log transformed data due to heteroskedasticity. Either two-tailed paired *t*-tests or one-, two-, or three-way mixed model ANOVAs (random factor: animal) were performed depending on the experimental design. Statistically significant *F* values (*p*<0.05) were followed by Tukey’s post hoc tests. Figures were created in Graphpad 9.4.1. Within-group individual differences were evident, thus for ease of visual interpretation we depict data that is normalized to a within-animal condition described in each figure legend. Data analysis was performed on raw, non-normalized data.

## RESULTS

### Region and culture density, but not morphine exposure, influence the survival of microglia *in vitro*

Our goal was to conduct a four-variable study directly comparing sex, region, density, and morphine concentration on microglial phagocytic outcomes. First, to ensure the selected morphine concentrations would not increase cell death, we measured live cells prior to plating versus at the conclusion of the 3.5 hr experimental protocol (2 hrs acclimation in media, 1.5 hrs morphine exposure; **Fig. 2**). Data were analyzed with a three-way ANOVA (drug x region x density). We unexpectedly found that while morphine did not impact cell survival (main effect of drug: *F*_3,41_=0.692, *p*=0.562), there were main effects of both region and density (region: *F*_1,41_=4.608, *p*=0.038; density: *F*_1,41_=124.885, *p*<0.001). Specifically, microglia from striatum survived better than microglia from cortex, and microglia plated at 50K densities survived better than microglia plated at 10K densities. There was no significant interaction between variables in cell survival. Although we were not powered for sex in these studies, we did perform the four-way ANOVA adding sex as a variable to ensure we were not missing a large effect, which was indeed absent (*F*_1,2.013_=0.390, *p*=0.596).

**Figure 2.**
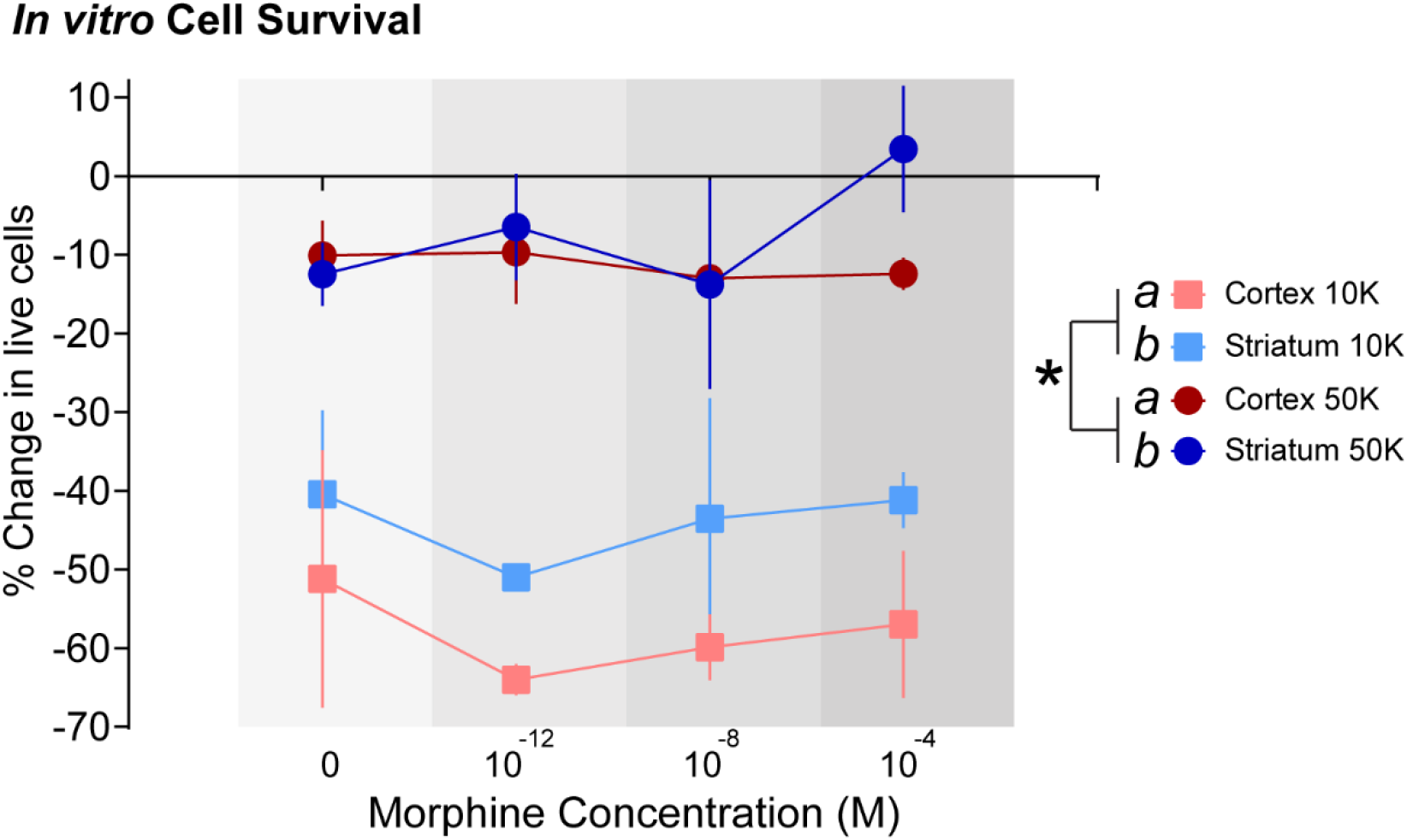
Region and culture density, but not morphine exposure, influence the survival of microglia *in vitro*. The percent change in Trypan-Blue indicated live cells from pre-plating to post-experiment was calculated for striatal and cortical microglia plated at 10K or 50K densities and exposed to different concentrations of morphine for 90 mins. *n* = 3-4 rats (sexes combined). Circles depict average, vertical lines depict standard error of the mean. A three-way mixed model ANOVA revealed region and density, but not drug or interactions between variables, impact microglial survival. * indicates the main effect of density (50K > 10K). *a* and *b* indicate the main effect of region (Striatum > Cortex). Main effects were considered significant if *p*<0.05.

### 10^-12^M morphine increases phagocytic activity in cortical microglia plated at 50K, but not 10K cells per well, and in striatal microglia at either density, independent of sex

We could not directly compare cortex and striatum, or 10K and 50K densities due to their differential regulation of microglial survival *in vitro* (**Fig. 2**). Thus, we next sought to determine whether sex or morphine concentration would play a role in phagocytic activity in 10K cortex, 10K striatum, 50K cortex, and 50K striatum conditions. Microglia in each condition, and from male or female rats, were concurrently incubated with fluorescent microbeads and either 0M, 10^-12M^, 10^-8^M, or 10^-4^M concentrations of morphine for 1.5 hrs after the 2hr media acclimation. Data were analyzed with a two-way ANOVA (drug x sex). There was a main effect of drug in 10K striatum (*F*_3,37_=3.550, *p*=0.024), 50K cortex (*F*_3,38_=3.481, *p*=0.025), and 50K striatum groups (*F*_3,39_=3.102, *p*=0.038), but not in 10K cortex groups (*F*_3,33_=1.325, *p*=0.283; **Fig. 3**). Specifically, 10^-12^M morphine increased phagocytic activity relative to all other concentrations in 10K Striatum and 50K striatum groups and increased phagocytic activity relative to 0M and 10^-4^M morphine in 50K cortex groups. There was no significant effect of sex, nor a significant interaction between sex and drug concentration, in any condition.

**Figure 3:**
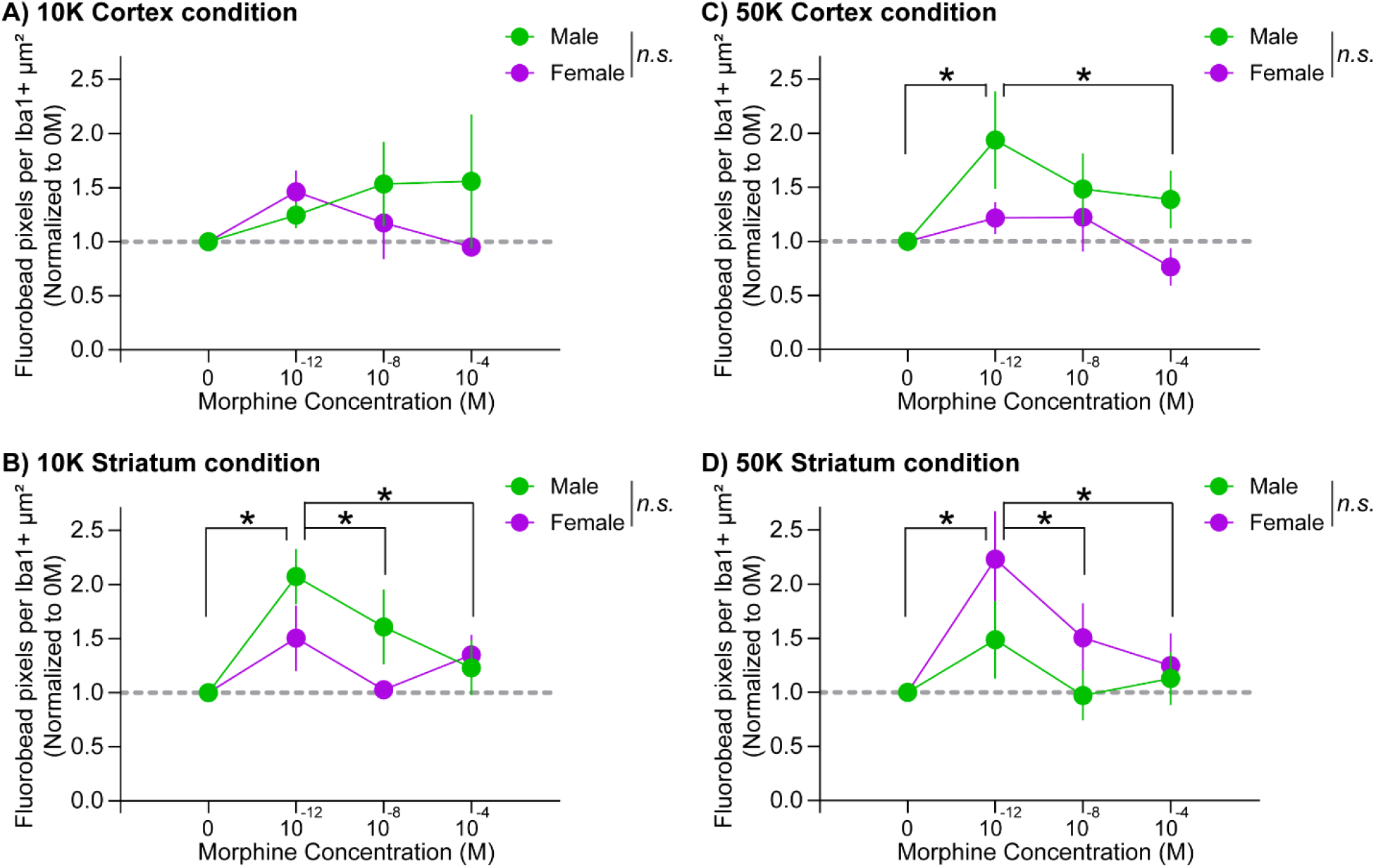
10^-12^M morphine increases phagocytic activity in cortical microglia plated at 50K, but not 10K cells per well, and in striatal microglia at either density, independent of sex. **(A)** There was no effect of morphine in cortical microglia plated at 10K density, while **(B)** 10^-12^M morphine increased phagocytic activity relative to all other concentrations in striatal microglia plated at 10K density. 10^-12^M morphine also increased phagocytic activity in both **(C)** cortical and **(D)** striatal microglia plated at 50K density (cortical: relative to 0M and 10^-4^M, but not 10^-8^M concentrations; striatal: relative to all other concentrations). Sex did not regulate phagocytic activity independently or via interactions with morphine concentration in any context. Circles depict mean, vertical lines depict standard error of the mean. Data were normalized to within-animal 0M conditions for ease of visualization. Data above the gray dotted line would indicate an increase in phagocytic activity above 0M control levels. * = *p*<0.05; *n* = 6-8 rats/sex.

### Morphine-induced increase in phagocytic activity in striatal microglia plated at 10K density is TLR4-, but not µOR-dependent

We next wanted to determine whether 10^-12^M morphine was increasing phagocytosis via TLR4- or µOR-dependent mechanisms. Pilot studies using DAMGO as an opioid receptor agonist and naltrexone as a µOR antagonist indicated that 100µM naltrexone would be effective in precluding µOR-dependent changes in microglial phagocytic activity (**Supplemental Fig. 2A**). Pilot studies using LPS as a TLR4 agonist and RS-LPS as a TLR4 antagonist indicated that 1µg/mL RS-LPS would be effective in precluding TLR4-dependent changes in microglial phagocytic activity (**Supplemental Fig. 2B**). Thus, we incubated cortical and striatal microglia plated at 10K densities with either 100µM naltrexone (or vehicle), 1µg/mL RS-LPS (or vehicle), the combined inhibitors, or combined vehicles for 30 mins. Then, we added a solution containing Fluorescent microbeads, 10^-12^M morphine (final concentration), and the respective inhibitors to maintain their final concentrations, for an additional 90 mins. There were 8 total treatment groups, and because we found no significant sex difference in the effects of morphine on phagocytosis in the studies depicted in **Fig. 3**, we combined the data from male and female rats. One-way ANOVAs indicated there was no significant difference between Vehicle conditions in either 10K Cortex (*F*_3,42_=1.475, *p*=0.235) or 10K Striatum (*F*_3,43_=2.382, *p*=0.083) microglia, and thus we used a single averaged vehicle value for each animal for data analysis. Paired *t*-tests from within-animal vehicle and 10^-12^M morphine + inhibitor vehicle data indicated that the data in **Fig. 3** did replicate: we found no significant effect of morphine on phagocytosis in 10K Cortex conditions (*t*(14)=1.669, *p*=0.117), but morphine increased phagocytosis in 10K Striatum conditions (*t*(15)=2.301, *p*=0.0362). A one-way ANOVA indicated there was an effect of inhibitor on morphine-induced phagocytosis (*F*_3,43_=7.222, *p*<0.001). Subsequent Tukey’s posthoc tests indicated the morphine-induced increase in phagocytosis in 10K Striatum microglia was inhibited by RS-LPS (*p*=0.002) or RS-LPS+naltrexone (*p*<0.001), but not by naltrexone alone (*p*=0.204). To confirm that sex did not play a role in these data, we also performed a two-way ANOVA (inhibitor x sex), and sex was not found to play a statistically significant role independently (*F*_1,14.026_=3.085, *p*=0.101) or via interaction with inhibitors (*F*_3,40_=2.066, *p*=0.120).

## DISCUSSION

Herein, we demonstrate that the brain region from which microglia are isolated and plating density both impact cell survival *in vitro* (**Fig. 2**). We found that 10^-12^M morphine, but not higher concentrations, increases phagocytosis in striatal microglia *in vitro* independent of sex and plating density (**Fig. 3B, D**). 10^-12^M morphine also increased phagocytosis in cortical microglia *in vitro* independent of sex, but this was contingent on a plating density of 50K (versus 10K; **Fig. 3A, C**). This effect was reliable, as an independent replication of the effect of 10^-12^M morphine on 10K cortical and striatal microglia also showed no change in phagocytosis, and increased phagocytosis, respectively (**Fig. 4A**). Finally, we demonstrate that the effect of 10^-12^M morphine in 10K striatal microglia is mediated via TLR4, and not µORs (**Fig. 4B**).

**Figure 4:**
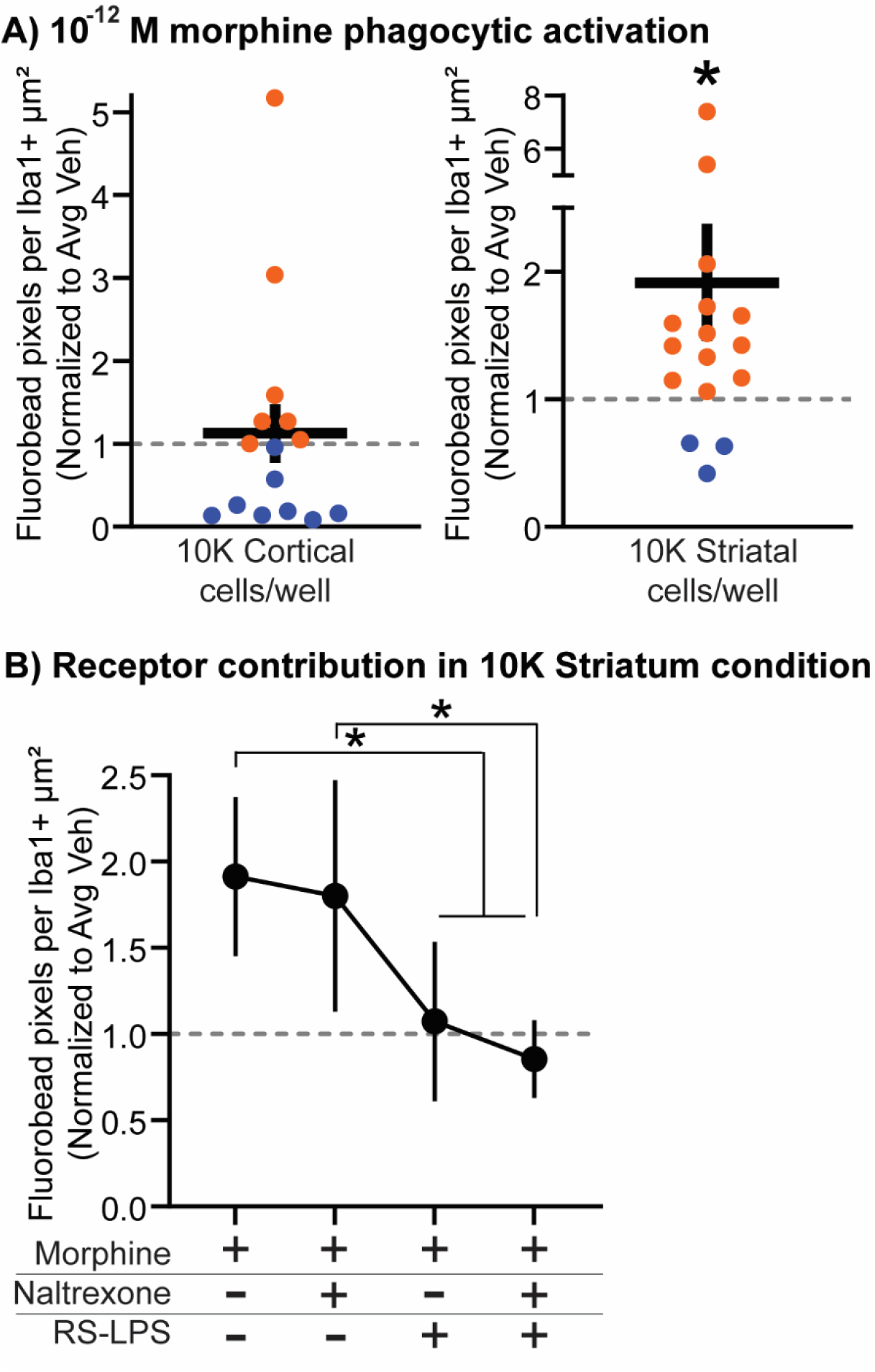
Morphine-induced increase in phagocytic activity in striatal microglia plated at 10K density is TLR4-, but not µOR-dependent. (A) A separate set of studies replicated the increased phagocytic effect of 10^-12^M morphine on striatal, but not cortical microglia plated at 10K density. Data are normalized to within-animal vehicle conditions for ease of visualization. Data above the gray dotted line would indicate an increase in phagocytic activity above 0M control levels and are identified with orange circles; data below the gray dotted line would indicate a decrease in phagocytic activity and are identified with blue circles. * = *p*<0.05; *n* = 15-16 rats (sexes combined). (B) Inhibiting TLR4s with RS-LPS, or TLR4s and µORs with a combination of RS-LPS and naltrexone, but not µORs (naltrexone) alone, inhibited the effect of 10^-12^M morphine on phagocytosis in striatal microglia plated at 10K density. One morphine+naltrexone+RS-LPS-sample was identified as an outlier and removed from analysis. Data were normalized to within-animal vehicle conditions for ease of visualization. Data above the gray dotted line would indicate an increase in phagocytic activity above 0M control levels. Circles depict mean, vertical lines depict standard error of the mean. * = *p*<0.05; *n* = 15-16 rats (sexes combined).

We initially set out to directly compare the effects of culture density, region, sex, and morphine concentration, as well as their interactions, on microglial phagocytic activity. However, several unexpected findings limited our ability to directly compare the effects of these variables concurrently. We predicted that a higher culture density (50K) would increase phagocytic activity relative to lower (10K) culture density. However, we found that microglia plated at 10K densities did not survive as well as microglia plated at 50K densities, and thus it was not appropriate to make this direct statistical comparison (**Fig. 2**). This is important to consider when reconciling the literature, which include highly variable culture densities with density rarely included as a variable. In two reports in which microglial phagocytosis is increased by morphine, the plating density was not reported in one (as far as we could find) and was 30,000 cells/well in chambered slides (Lipovsky et al., 1998; Peterson et al., 1995). In two reports in which microglial phagocytosis was decreased by morphine, the plating density was 50,000 cells/ml and 40,000 cells/well (Ryu et al., 2018; G. Sowa et al., 1997). Although we observed that morphine generally increased phagocytosis in 50K and 10K conditions, this did not reach statistical significance in cortical microglia plated at 10K density. It is thus possible that cell viability, perhaps in interaction with region examined, may play a role in these disparate conclusions. Finally, although we used an artificially dramatic difference in microglial density to test our hypothesis, it is worth noting that microglia density does vary across the brain, particularly in the substantia nigra which has the highest density of microglia (Lawson et al., 1990; Mittelbronn et al., 2001; Wang et al., 2015). Our data might suggest that a higher density of microglia might be protective in terms of cell viability, but there are several reports indicating that substantia nigra might be a particularly vulnerable region *because* of its high density of microglia (Flores-Martinez et al., 2018; Milde et al., 2021; Wang et al., 2014). There is likely to be a difference between microglial density in acute (e.g., in our experimental design) vs. chronic conditions. It may be worth varying culture density in long-term cultures to test whether lower microglial density has a negative effect under chronic activation or inflammatory conditions.

We predicted that striatal microglia would be more phagocytically active than cortical microglia. Activation of microglia under normal and pathological states increases pro-inflammatory profiles in striatum more than in cortex (Crapser et al., 2020; Klawonn et al., 2021). We observed that striatal microglia survived better than cortical microglia, and thus a direct comparison was not possible (**Fig. 2**). Although this was a main effect, not an interaction between density and region, visual inspection of the data indicate that this effect is driven primarily by worse survival outcomes in cortical microglia plated at 10K density. We propose that the beneficial effects of higher density culture conditions are a result of inter-cellular signaling promoting survival. If this is the case, cortical microglia may be particularly vulnerable (or striatal microglia more resilient) to the presence or absence of these signals. There are currently no direct comparisons of cell survival or death signaling profiles between cortical and striatal microglia. However, microglial apoptotic regulation is linked to reduced phagocytosis and reduced responses to viral load (Jeong et al., 2022; Liu et al., 2001). Thus, a few theories might explain the fact that morphine did not increase phagocytosis in 10K cortical microglia (**Fig. 3A**): (1) reduced survival might bias our current data, (2) activated apoptotic signaling might inhibit phagocytic potential, and (3) morphine does not affect microglia in this context. In future studies it will be worthwhile to examine higher density microglia from different regions to remove the possibility of differential survival.

We predicted that microglia from female rats would be more phagocytically active than microglia from male rats. To our surprise, we did not observe any significant effects of sex, neither main effects nor interactions, in any of our experiments (**Fig. 3**). This may be due to the rats being young adults, past the developmental plasticity that might be driving sex differences in microglia (Lynch, 2022; Osborne et al., 2019), and prior to when aging might again cause sex differences in microglia (Daly et al., 2022; Mela et al., 2022). If true, this raises the question: what do embryonic or immortalized cultures represent? The general consensus is that these longer-term culturing methods do not reflect mature cell phenotypes (He et al., 2018; Stansley et al., 2012). That may be a distinct disadvantage depending on the questions being asked, just as our current design is unlikely to be reflective of microglial responses at a different life stage. The lack of a finding of a sex difference may also be due to being underpowered to detect subtle effects of sex in a complex experimental design. For example, there may be sex x drug concentration interactions in the 50K Cortex condition that we cannot resolve with the effect sizes we attained (**Fig. 3C**). Finally, other pharmacological activators of phagocytosis such as lipopolysaccharide result in sex differences (Loram et al., 2012; Thion et al., 2018), and so does morphine in peripheral immune cells (Doyle & Murphy, 2017; Rosen et al., 2019). There are reported sex differences in the response to opioids in humans and rodent models (Bobzean et al., 2019; Boorman & Keay, 2022; Escorial et al., 2022), but microglial phagocytosis need not mediate those effects.

We hypothesized that morphine would decrease phagocytosis in a dose-dependent manner. Morphine penetrance into the brain is dependent on concentration, route of administration, metabolism, and inter-individual factors (Chaves et al., 2017; Ollikainen et al., 2019; Rogers et al., 2013; Smith, 2009). The statistically significant main effects indicate that 10^-12^M morphine increased phagocytosis in 10K Striatum, 50K Cortex, and 50K Striatum conditions (**Fig. 3B-D**). There was no significant effect of morphine in 10K Cortex conditions (**Fig. 3A**). It is possible that lower survival in this condition is resulting to the null effect, but visual inspection of the data indicates that story is likely to be more complicated. While there is no statistically significant effect of morphine on cortical microglia plated at 10K density, there is a rather clear pattern of 10^-12^M morphine increasing phagocytosis relative to other concentrations – in females only. This effect is likely masked (from a statistical analysis perspective) by high variance in male data that does not follow the same trend. Larger group sizes may permit more nuanced conclusions. We find that at lower morphine concentrations phagocytosis is increased. This suggests that there is a window of increased microglia phagocytosis *in vivo* which could be immediate or delayed based on peak brain morphine concentration as the concentration decays with time. Importantly, whether there is a sex difference in the effect of morphine on microglial phagocytosis or not, it does appear that morphine is generally increasing phagocytosis. This could be consequential for the effect of morphine on synaptic density, as synapses are a frequent target of microglial phagocytosis (Authement et al., 2016; Kopec et al., 2018; Zhang et al., 2021). In fact, there is reports that synaptic density is decreased in nucleus accumbens and intermediate medial interstitial after morphine exposures (Kasture et al., 2009; Wang et al., 2023). An open question is whether microglia would be a mediator of that effect.

Morphine is documented to activate both TLR4 and µOR signaling to mediate its effects. Thus, in our final set of studies, we tested whether increased phagocytosis in 10^-12^M morphine treated striatal microglia plated at 10K density required TLR4s, µORs, or both. The data indicate that TLR4, but not µORs, is important for this effect (**Fig. 4**). While the expression of TLR4 is enriched in microglia in neural cell datasets, there has been some controversy over whether microglia express µORs. The morphine binding affinity for µOR (Human) has been measured at ∼1.14nM (Volpe et al., 2011) and morphine levels in the brain peak in the 10-500 ng/ml range depending on the morphine metabolite measured (Barjavel et al., 1994; Barjavel et al., 1995; Ederoth et al., 2004; Stain-Texier et al., 1999). However, morphine has been shown to modulate microglia in femtomolar (10^-15^) ranges (Grzegorz Sowa et al., 1997). While morphine has been shown to bind to TLR4 in a parallel manner to its well-known agonist endotoxin (lipopolysaccharide, specifically) (X. Wang et al., 2012), the binding affinity of morphine for TLR4 is an ongoing endeavor (J. H. L. Thomas et al., 2022). However, in peripheral macrophages TLR4 potentiates phagocytosis more robustly than opioid receptors (Ninkovic et al., 2016) and sex-specifically influences antinociception, tolerance and withdrawal (Jacob H. L. Thomas et al., 2022). Both opioid receptors and TLR4s are reported to influence the effects of morphine on microglial immune signaling (Liang et al., 2016; Lipovsky et al., 1998; X. Wang et al., 2012; Xie et al., 2017; Y. Yang et al., 2021). Our pilot data indicate that microglia respond to a non-specific opioid receptor agonist (DAMGO) with increased phagocytosis, and a µOR-specific antagonist (naltrexone) can inhibit this effect. These data would argue that µORs are, in fact, expressed and functional in microglia. However, this was a pilot study and underpowered for making conclusive statements. We thus cannot rule out that the null effect of naltrexone on morphine-induced phagocytosis is due to no or low expression of µORs on microglia. Regardless, our data do indicate that morphine is activating microglia through canonical immune pathways. TLR4 signaling is associated not just with phagocytic regulation, but also pro-inflammatory cytokine induction. Indeed, morphine potentiates synthesis and release of IL-1β and TNF-α in microglia (Liang et al., 2016; Y. Yang et al., 2021). Whether this would be the case in microglia from either region plated at 50K remains to be tested. It is possible that a µOR- and region-dependent effect would emerge in microglia plated at 50K due to their greater survival (and thus wider sampling), because striatum contains microglia exposed to higher endogenous opioidergic signaling (Rocchi et al., 2020).

### Limitations to the current studies

The choice of experimental conditions in studies such as these is not a trivial matter (Bohlen et al., 2017). We made several experimental decisions that could impact our conclusions. Some of the major caveats include: (**1**) Our data comes from microglia from a single rat strain (Sprague-Dawley) and further studies on other rat strains and animal species are needed for generalization of these results. (**2**) Media plays an enormous role in *in vitro* outcomes. Our media does not contain serum, which is known to impact microglial health in culture, but does contain low-concentration (1µM) forskolin. We used forskolin to upregulate cAMP levels in microglia, as cAMP–regulated pathways are essential for physiological cell processes (Sassone-Corsi, 2012) which are stunted during cell isolation (Bohlen et al., 2019). Additionally, cAMP pathways are important for pro-phagocytic responses such as deramification and surveillance, hallmarks of phagocytic activity (Bernier et al., 2019; Ghosh et al., 2020). However, forskolin has been linked to both enhancement and suppression of neuroinflammation (Owona et al., 2016; Veremeyko et al., 2018; Woo et al., 2004). Opioid receptor agonists such as morphine inhibit the generation of cAMP (Zhang et al., 2020), and upregulating cAMP is protective against morphine-induced microglia consequences (He & Whistler, 2005). The 1µM concentration of forskolin used herein is lower than typical studies using forskolin (often 10µM), and there are clear phagocytic responses to morphine in striatum and to DAMGO and LPS in whole brain. Finally (**3**), any isolation protocol will impact the cells in a non-physiological way. We use dissociation techniques reported to perform well in terms of microglial health and viability (Bordt et al., 2020; Nikodemova & Watters, 2012), but there is no way to attain a true *in vivo* physiological setting *in vitro*.

## CONCLUSIONS

Overall, our data suggest that a morphine-TLR4 signaling pathway increases phagocytic activity in microglia independent of sex, which is useful information for better understanding the possible neural outcomes associated with morphine exposures.

## Acknowledgements

We thank Dr. Justin Bourgeois and Erin Edgar for experimental and quantification assistance. We thank Dr. Margarida Barroso and the AMC Imaging Core Facility for imaging assistance.

## Author Declarations

DNK, LN, JLB, KCM, and AMK designed the experiments. DNK, LN, JLB, and AMK performed the experiments. DNK, LN, JLB, PJF, and AMK analyzed the experiments. DNK, MT, and AMK wrote the manuscript. All authors edited the manuscript. The authors declare no conflicts of interest.

## FIGURES

**Supp. Figure 1:**
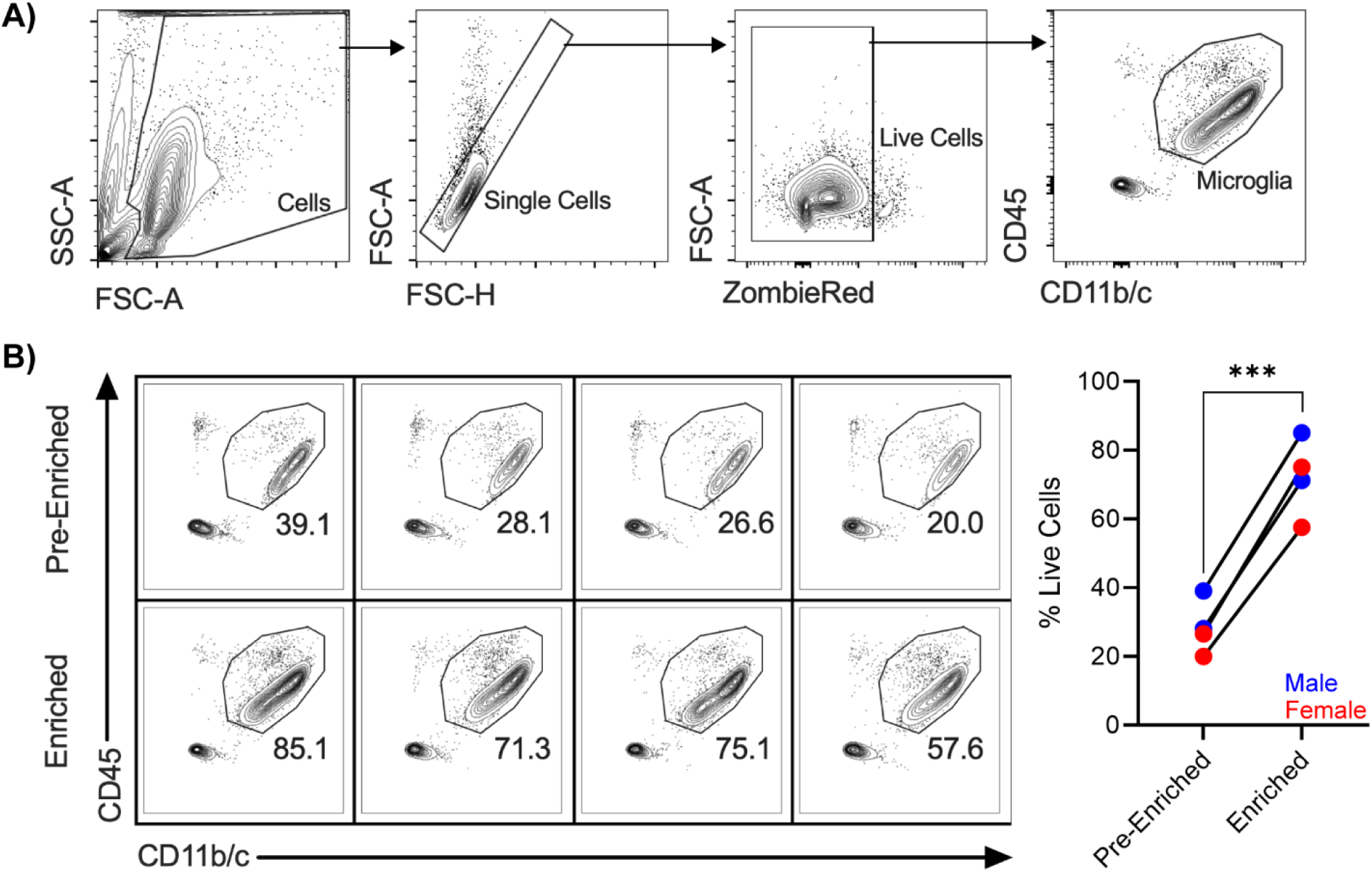
CD11b magnetic isolation enriches for microglia. Microglia were enriched from single-cell suspension of rat brain tissue using Miltenyi Biotec MicroBeads. **(A)** Cells were stained with antibodies against CD11b/c and CD45 and analyzed via flow cytometry. **(B)** The frequency of microglia was compared before and after magnetic microbead enrichment. Data were analyzed with a paired *t*-test. *** = p<0.001; *n*=4 rats (sexes combined).

**Supp. Figure 2:**
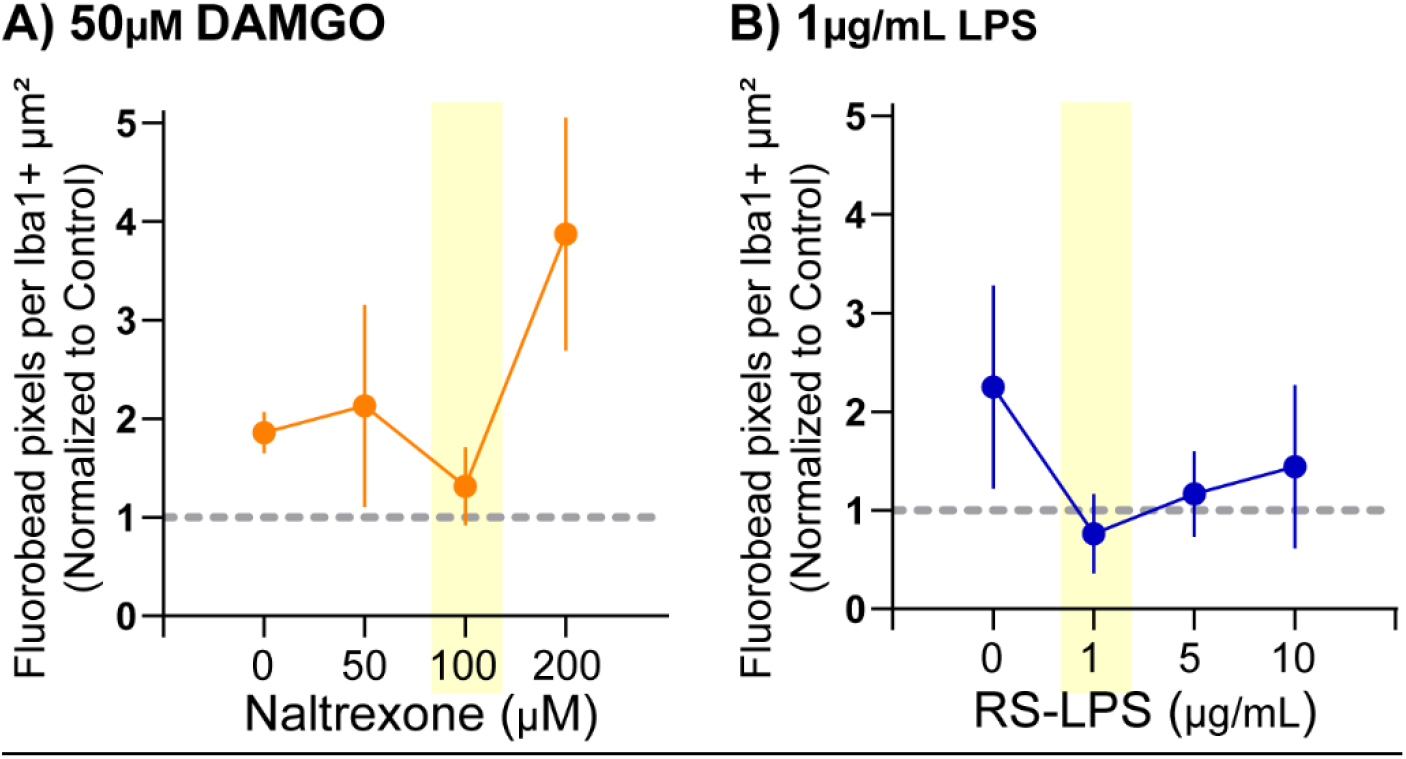
100µM naltrexone and 1µg/mL RS-LPS reduce DAMGO- and LPS-induced phagocytic activity, respectively. **(A)** Microglia from whole brain were plated at 50K density and exposed to fluorescent microbeads, Vehicle or 50µM DAMGO, an opioid receptor agonist, and varying concentrations of naltrexone, a µOR antagonist. 100µM naltrexone reduced phagocytosis induced by DAMGO to control levels. **(B)** Microglia from whole brain were plated at 50K density and exposed to fluorescent microbeads, Vehicle or 1µg/mL LPS, a TLR4 agonist, and varying concentrations of RS-LPS, a TLR4 antagonist. 1µg/mL RS-LPS reduced phagocytosis induced by LPS to control levels. Data were normalized to within-animal vehicle conditions (an average of 0M agonist + each inhibitor concentration) for ease of visualization. Data above the gray dotted line would indicate an increase in phagocytic activity above control levels. Circles depict mean, vertical lines depict standard error of the mean. *n* = 3-4 rats (sexes combined). Statistics were not performed as this was a pilot experiment to choose inhibitor concentrations for the experiment displayed in Figure 5.

